# Evaluating Nanopore read trimming and quality filtering in bacterial genomics

**DOI:** 10.64898/2026.10.12.758701

**Authors:** Achmad Dimas Cahyaning Furqon, Ryan R Wick, Leah W Roberts, Michael B Hall

**Author notes:** These authors share senior authorship (Leah W Roberts, Michael B Hall).

## Abstract

Oxford Nanopore Technologies (ONT) sequencing is widely used in bacterial genomics, particularly for reference-based variant calling and *de novo* assembly. Both analyses routinely start with quality control (QC) preprocessing, but as ONT basecalling accuracy has improved, its necessity and downstream trade-offs remain poorly defined. We benchmarked 17 preprocessing combinations, built from three adapter trimmers (Dorado, Porechop ABI, Barbell) and three quality-based processing tools (Filtlong, Chopper, SeqKit) with untrimmed and unprocessed controls, plus Fastplong, which performs both. Each combination was evaluated across 13 bacterial isolates at 100× and 20× depth under the HAC and SUP basecalling models. Variant calling was assessed by F1 score. Assemblies were scored on sequence accuracy, contiguity, replicon recovery and residual adapter contamination, then aggregated into a weighted arithmetic mean using weights from a survey of the microbial genomics community. With SUP basecalling, raw reads with no trimming or quality-based processing produced accurate SNP calls and outperformed several pipelines that applied both operations; adapter trimming alone left SNP accuracy unchanged. Subsetting reads by combined length and quality with Filtlong gave the highest SNP F1 scores at both depths, except with Barbell at 20×, where it ranked second lowest. As is often the case in benchmarking studies, assembly results involved trade-offs, and the best tool depends on which aspects of performance matter most for a given use. Filtlong produced the lowest error rates, but every Filtlong pipeline lost three small plasmids at both depths. Barbell trimming raised error rates and reduced contiguity at 20×, while skipping adapter trimming left residual adapter and barcode sequences in assemblies. Under the community-derived weights, Porechop ABI followed by Chopper had the highest overall score at both depths, without winning any single metric. QC tool choice should therefore be guided by the downstream analysis goal rather than applied indiscriminately across all workflows.

**Author summary:** Oxford Nanopore reads are routinely processed to remove adapter and barcode sequences introduced during library preparation and discard low-quality reads or read segments. As sequencing accuracy improves, the benefits and costs of these steps need reassessment. We compared 17 preprocessing combinations across 13 bacterial isolates for detecting genetic differences (variant calling) and reconstructing genomes (genome assembly). Most combinations produced similar results to unprocessed reads, but some affected the genetic information recovered. Filtlong, which selects reads by length and quality, improved detection of single-nucleotide variants in several combinations and reduced assembly errors. However, it removed reads supporting three small plasmids, preventing their assembly. Missing small plasmids in bacterial assemblies can have a large impact, as they are known to carry important genes (e.g., antibiotic resistance). Adapter trimming offered little benefit for single-nucleotide variant detection, but omitting it left adapter and barcode sequences in some assemblies. These findings support choosing preprocessing tools according to the intended analysis. We surveyed researchers to weight four aspects of assembly quality: sequence accuracy, contiguity, chromosome and plasmid recovery, and residual adapter contamination. Our interactive dashboard lets users adjust these weights to compare pipelines according to their own priorities.

## Introduction

Oxford Nanopore Technologies (ONT) sequencing is now widely used in clinical microbiology, evolution, and epidemiological surveillance. One major application is variant calling, in which reads are aligned to a reference genome and single nucleotide polymorphisms (SNPs), insertions, deletions (indels), and structural variants are identified. These variants matter for resistance-associated mutations, because SNPs or indels in coding or regulatory regions can alter gene expression or function [1, 2], and SNPs are also used to classify lineages through phylogenetic approaches and to inform epidemiological surveillance [3, 4]. The other major application is *de novo* genome assembly, where genomes are reconstructed directly from the reads without a reference. This reduces reference bias and preserves genetic context: the position, orientation, and arrangement of genes and mobile elements [5, 6].

Quality control (QC) preprocessing is a standard step before both workflows, with adapter trimming and quality-based processing being the most common operations. Trimming removes unwanted sequence segments from a read, such as residual adapter, barcode, and primer sequences at the read ends or within the read [7, 8]. Quality-based processing removes either entire reads (filtering) or sections of a read that do not meet a quality or length threshold [9]. Some filters rank reads by a combination of length and quality and keep only the highest-ranked subset up to a target number of bases [10]. What a trimmer has to remove depends on how the library was prepared: ligation-based native barcoding attaches barcode and adapter sequence at both ends of a fragment, whereas tagmentation-based rapid barcoding attaches them at one end [11, 12, 13].

Whether these preprocessing steps provide sufficient benefit to justify their associated loss of data is not yet clear. The closest evidence comes from short-read sequencing. Bush compared several trimming tools in bacterial Illumina variant calling pipelines, and found that trimming changed relatively few calls, although it reduced mixed calls in some analyses [14]. At the same time, evidence from ONT studies remains mixed. In genome assembly, Beeloo *et al*. showed that barcode and adapter sequences can survive ONT demultiplexing and trimming, and then propagate into assemblies and public sequence records [13]. Other work, however, found that long-read assembly was largely unaffected by adapter sequences and other problematic reads, including random, junk, and chimeric reads [15]. At the same time, improvements in ONT basecalling accuracy [16, 17], together with advances in variant calling models and long-read assembly, have improved the accuracy of downstream analyses [17, 18, 19]. These developments raise the question of whether preprocessing steps developed to address earlier sequencing limitations remain necessary.

The effect of preprocessing is also unlikely to be the same for every downstream analysis or dataset. Assembly and variant calling do not ask the same thing of the reads that are kept [17, 20], so one preprocessing choice can improve one outcome while costing another, which a benchmark that reports only a single summary number will not show. Assembly and variant calling performance also depend on sequencing depth and basecalling accuracy [15, 17], so the effect of preprocessing may depend on the depth and basecalling model of the data being processed. No existing study compares adapter trimming and quality-based processing tools jointly across basecalling models, sequencing depths, and both downstream workflows.

In this study, we benchmarked 17 combinations of adapter trimming and quality-based processing, including a raw control that applied neither, across 13 bacterial isolates, each from a different species, sequenced with two ONT barcoding kits. We evaluated every combination at two sequencing depths and under two basecalling models, for reference-based variant calling and for *de novo* genome assembly. Variant calling performance was assessed primarily using F1 score, with recall, precision, and the numbers of false negative and false positive calls used as supporting metrics. Assemblies were assessed on four criteria: sequence accuracy, contiguity, replicon recovery, and absence of residual adapter and barcode sequence. We combined these assembly metrics into a single score using weights extracted from a survey of the microbial genomics community. Because those weights describe only one community sample, rather than the priorities of any particular project, we also provide an interactive dashboard that allows a reader to reweight the criteria and apply their own thresholds.

## Materials and methods

### Dataset and basecalling

Raw ONT signal data (POD5 files) were obtained from Hall *et al*. [17]. This dataset consists of 13 bacterial species sequenced on either a MinION Mk1b or GridION device using ONT R10.4.1 flow cells (FLO-MIN114), along with their corresponding ground-truth reference assemblies, mutated references, and variant truthsets [21]. For each isolate, Hall *et al*. identified the SNPs and indels (up to 50 bp) that distinguish its reference genome from a closely related donor genome (98.4–99.8% average nucleotide identity), and applied these variants to the reference to produce a mutated reference [17]. Reads from the isolate therefore differ from its mutated reference at exactly these sites, and these differences form the variant truthset for that isolate.

To avoid confounding adapter and barcode detection, each sample comprised POD5 files from a single sequencing kit. An initial 14th sample (KPC2202310) was excluded because its single-kit sequencing depth fell below the 100× target. The final dataset was balanced across kits, with seven samples sequenced using the Native Barcoding Kit V14 (SQK-NBD114-96) and six using the Rapid Barcoding Kit V14 (SQK-RBK114-96), as summarised in Table 1.

**Table 1.** Dataset composition. Overview of the 13 bacterial samples evaluated in the study. Sequencing kits are SQK-NBD114-96 (Native Barcoding Kit) and SQK-RBK114-96 (Rapid Barcoding Kit). Read counts and total yield reflect the raw reads, before adapter trimming or quality-based processing, from the SUP basecalling model.

| Sample | Species | Sequencing Kit | Read Count | Yield (Mbp) |
| --- | --- | --- | --- | --- |
| ATCC_10708__202309 | *Salmonella enterica* | SQK-RBK114-96 | 843,884 | 2,247.3 |
| ATCC_17802__202309 | *Vibrio parahaemolyticus* | SQK-RBK114-96 | 568,955 | 1,642.4 |
| ATCC_19119__202309 | *Listeria ivanovii* | SQK-RBK114-96 | 671,362 | 2,388.3 |
| ATCC_25922__202309 | *Escherichia coli* | SQK-RBK114-96 | 1,093,172 | 4,017.2 |
| ATCC_35897__202309 | *Listeria welshimeri* | SQK-RBK114-96 | 1,022,948 | 3,876.9 |
| ATCC_BAA-679__202309 | *Listeria monocytogenes* | SQK-RBK114-96 | 911,644 | 2,794.8 |
| ATCC_33560__202309 | *Campylobacter jejuni* | SQK-NBD114-96 | 564,305 | 1,806.7 |
| ATCC_35221__202309 | *Campylobacter lari* | SQK-NBD114-96 | 443,217 | 1,592.6 |
| AJ292__202310 | *Klebsiella variicola* | SQK-NBD114-96 | 71,992 | 587.4 |
| RDH275__202311 | *Streptococcus pyogenes* | SQK-NBD114-96 | 1,013,240 | 2,862.9 |
| MMC234__202311 | *Streptococcus dysgalactiae* | SQK-NBD114-96 | 236,637 | 970.4 |
| BPH2947__202310 | *Staphylococcus aureus* | SQK-NBD114-96 | 162,939 | 694.2 |
| AMtb_1__202402 | *Mycobacterium tuberculosis* | SQK-NBD114-96 | 187,246 | 719.2 |

POD5 files were basecalled with Dorado v1.4.0 using both the Super Accurate (SUP; dna_r10.4.1_e8.2_400bps_sup@v5.2.0) and High Accuracy (HAC; dna_r10.4.1_e8.2_400bps_hac@v5.2.0) models. Dorado was run both under default settings, which trim adapter and barcode sequences automatically, and with the `--no-trim` flag, which leaves those sequences intact.

### Read trimming and quality-based preprocessing

Fig 1 outlines the benchmarking workflow. Untrimmed FASTQ files retaining intact barcode and adapter sequences served as the negative control for adapter trimming, and as input for Barbell v0.3.2 [13], which was run in demultiplexing mode through the `barbell kit` subcommand under default parameters. FASTQ files from Dorado basecalling under default settings were processed with either Dorado v1.4.0 (`dorado trim`, default parameters) or Porechop ABI v0.5.0 [22] with the `--discard middle` flag to remove chimeric reads. Both Dorado and Barbell required specifying the corresponding sequencing kit for each sample (Table 1), whereas Porechop ABI identifies adapter sequences *ab initio* without requiring prior kit information.

**Fig 1.**
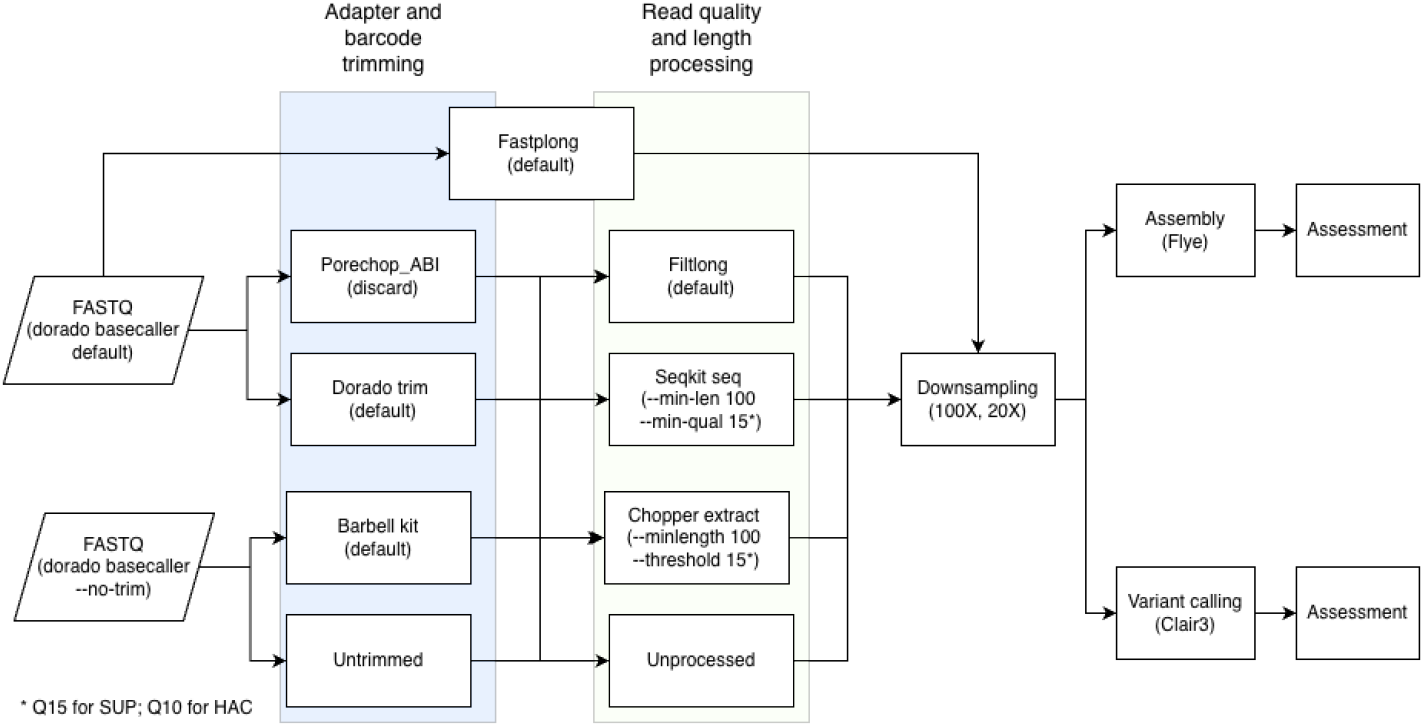
Overview of the benchmarking workflow. Raw ONT POD5 files were basecalled with Dorado using either default settings or the `--no-trim` option. Reads were processed using adapter and barcode trimming tools, followed by quality-based processing tools. Because Fastplong performs both operations simultaneously, it was evaluated as a standalone workflow. The resulting 17 preprocessed datasets were subsequently downsampled to target sequencing depths of 100× and 20×. Finally, these combinations were assessed for variant calling performance using Clair3 v2.0.1 and genome assembly performance using Flye v2.9.6.

Following adapter trimming, datasets underwent quality-based processing. This step evaluated three tools alongside an unprocessed control: Filtlong v0.3.1 [10] with default weighting parameters and a target base count of 100× the sample’s genome size; Chopper v0.12.0b [9] using the `best-read-segment` trimming approach; and SeqKit v2.13 [23]. For Chopper and SeqKit, the minimum read length was 100 bp, with minimum quality cutoffs of Q15 for SUP datasets and Q10 for HAC datasets. Tools offering several operational modes were assessed independently beforehand to select their parameters (Supplementary Section S1, Supplementary Figs S1–S4 and Supplementary Table S1).

This sequential workflow yielded 16 combinations of adapter trimming and quality-based processing tools. Fastplong v0.4.1 [7] performs adapter trimming without requiring a specified sequencing kit and quality filtering simultaneously, so it was evaluated as a standalone workflow using the length and quality thresholds applied to SeqKit and Chopper. With the untrimmed and unprocessed raw dataset as a negative control for both conditions, the benchmarking matrix totalled 17 preprocessing combinations. All processed datasets were then downsampled to target depths of 100× and 20× with Rasusa v3.0.0 [24].

### Variant calling

Processed reads from each combination were aligned to their mutated reference genomes with Minimap2 v2.30 [25] using the `map-ont` preset and the `-aL`, `--cs`, and `–MD` options, then coordinate-sorted with Samtools v1.23.1 [26]. Sorted BAM files were used for variant calling with Clair3 v2.0.1 [18], applying the `r1041 e82 400bps sup v520` and `r1041 e82 400bps hac v520` models to match the basecalling model.

Variant calling accuracy was benchmarked against the reference truthsets with VCFdist v2.6.4 [27]. Partial credit was disabled (`--credit-threshold 1.0`), evaluation was restricted to variants *≤* 50 bp (`--largest-variant 50`), and the maximum variant quality threshold (`--max-qual`) was set dynamically to the highest score in the assessed VCF. VCFdist computes performance across a continuous range of quality score cutoffs, so we standardised by reporting peak performance for each condition: for every combination of pipeline, depth, basecalling model and variant class, we identified the quality threshold that maximised the F1 score and extracted the precision, recall, F1 score, and absolute false negative (FN) and false positive (FP) counts at that threshold.

### Genome assembly

Processed reads were assembled with Flye v2.9.6 [28] using `--nano-hq`. Assembly quality was evaluated on four criteria: sequence accuracy, assembly contiguity, replicon recovery, and residual adapter and barcode contamination. All reference genomes and assembly contigs were rotated to a consistent biological origin (*dnaA, terL, repA*, or COG1474) using Dnaapler v1.4.0 [29]. This avoids artificial alignment breaks and penalty artefacts at sequence termini.

#### Adapter and barcode contamination

Contamination was defined as adapter or barcode sequences retained in the final assembly. Screening was restricted to the sequencing kit used for each isolate (Table 1). Contaminants were detected by aligning assembly contigs (query) against expected adapter and barcode sequences (target) with Minimap2 using short-read alignment parameters tuned to retain short matches without splicing (`-c -x sr -k 9 -w 1 -n 1 -m 14`).

Structural sequence arrangements vary by kit [12]: the Rapid Barcoding Kit has an internal barcode flanked by left and right adapter sequences, whereas the Native Barcoding Kit has distinct adapter and barcode/flank architectures [13]. Alignment coverage was evaluated across these sub-regions. A valid contamination hit required *≥* 90% sequence identity and *≥* 90% alignment coverage across at least one structural sub-region. Where several barcode alignments overlapped on one contig, the alignment with the highest dynamic programming score was retained. Each retained alignment was counted as one contaminant hit, so a single contig can carry more than one. For each hit we also recorded the contig’s circularity status as reported by Flye and the distance from the hit to the nearest contig end.

#### Replicon recovery

This metric measures whether a pipeline recovers all expected replicons, including chromosomes and small plasmids. Assemblies were aligned to their reference assemblies with Minimap2 using the `asm5` preset, and per-contig reference coverage was calculated with `samtools coverage`. A reference contig was classified as fully missed at 0% coverage and partially missed at *>* 0% and *<* 50% coverage.

#### Sequence accuracy

Reoriented assemblies were assessed against their complete reference genomes with QUAST v5.3.0 [30] using default parameters. Sequence accuracy was quantified as the frequency of base mismatches and indels per 100 kbp of aligned sequence, calculated by summing the mismatch and indel rates reported by QUAST.

#### Assembly contiguity

We assessed contiguity with auNGA, the area under the NGAx curve [31, 32, 33]. QUAST splits contigs at detected misassemblies and excludes unaligned sequence, leaving a set of aligned blocks [30]. auNGA summarises the length of these blocks relative to the reference and gives more weight to long blocks. Because a raw auNGA value depends on the size and number of replicons in the reference, we divided each assembly’s auNGA by that of its own reference value, in which every replicon is one block. This auNGA ratio is 1.0 when an assembly recovers each reference replicon once as a single aligned block. It falls below 1.0 with fragmentation and can rise above 1.0 with duplicated sequence. We report contiguity as the auNGA score, max(0, 1 *−* |auNGA ratio *−* 1|), so that 100% is best and deviation in either direction is penalised. The loss of a small plasmid barely changes the ratio, which is why replicon recovery was scored as a separate criterion. The formulas and worked examples are given in Supplementary Section S2.

### Assembly metric aggregation and community survey

To rank pipelines on assembly performance, we aggregated the four assembly metrics into a single score. Each metric was normalised to a 0–100 scale: contiguity by penalising deviation from an auNGA ratio of 1.0 symmetrically; sequence accuracy against a fixed policy anchor of 10.0 total error events per 100 kbp; contamination as the proportion of the 13 isolates with no detected adapter or barcode hits; and replicon recovery against a denominator of three missed replicons across the cohort, so that a pipeline recovering every replicon scores 100 and losing three or more scores 0.

Contiguity and sequence accuracy were scored per isolate and then averaged across the cohort, whereas contamination and replicon recovery were evaluated across the cohort as a whole. The normalisation formulas are given in Supplementary Section S3.

To set the weights, we asked the microbial genomics community to complete a survey on the perceived importance of the four assembly metrics. Each respondent distributed 100 points across the four assembly metrics according to perceived importance, and the mean response provided the baseline weights (Supplementary Table S2). The composite score is the weighted arithmetic mean of the four normalised metric scores.

## Results

The results below are for SUP-basecalled reads. The equivalent HAC figures are in the Supplementary material, and HAC results are summarised in the text where relevant.

### Which pipelines give the most accurate variant calls?

Across all 17 pipelines, mean SNP F1 scores for SUP-basecalled data fell within 0.09 percentage points of each other at 100× depth and within 0.22 percentage points at 20× (Fig 2A). The raw control, `Unprocessed-Untrimmed`, placed inside that range at both depths, with mean F1 scores of 99.97% at 100× and 99.86% at 20×, higher than several pipelines that applied both trimming and quality-based processing.

**Fig 2.**
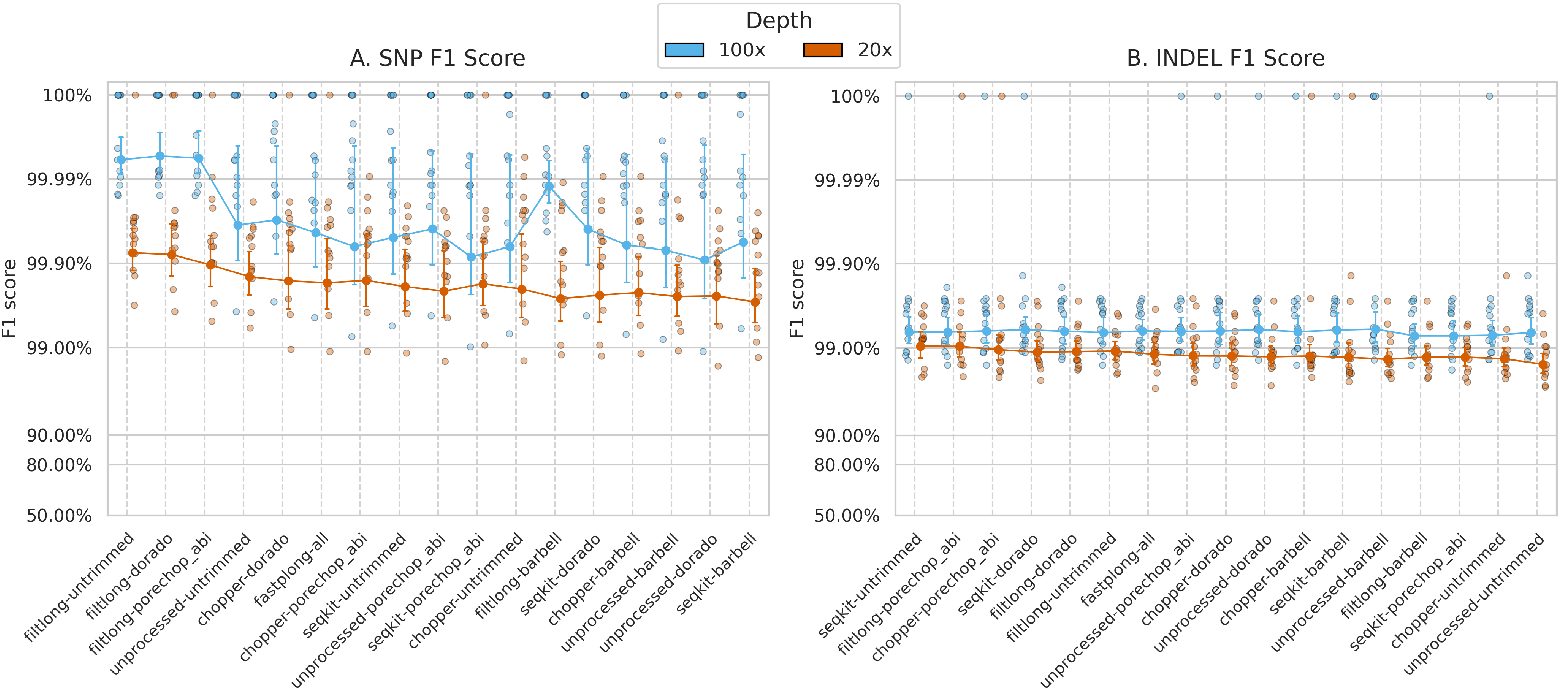
Variant calling performance across QC pipelines and sequencing depths. F1 scores for (A) SNP and (B) indel variant calling across the 17 pipelines at 100× (blue) and 20× (orange) depth, using SUP-basecalled reads. The y-axis is on a logit scale. Individual points are single-sample runs, solid markers are mean F1 scores, and error bars are 95% confidence intervals. QC pipelines along the x-axis are ordered from highest to lowest mean performance across both depths.

At 100×, the only three pipelines to exceed 99.99% all used Filtlong: `Filtlong-Dorado`, `Filtlong-Porechop ABI` and `Filtlong-Untrimmed`. At 20× the three highest means were again Filtlong pipelines, led by `Filtlong-Untrimmed` at 99.93%. The exception was `Filtlong-Barbell`, which performed close to the bottom of the range at 20× (99.74%), second lowest only to `SeqKit-Barbell` (99.71%). Adapter trimming on its own brought no gain: `Unprocessed-Dorado`, `Unprocessed-Porechop ABI` and `Unprocessed-Barbell` all fell slightly below the raw control, by at most 0.06 percentage points at 100× and 0.10 percentage points at 20×. HAC-basecalled data showed the same ordering, with all four Filtlong pipelines ranked highest at 100× (Supplementary Fig S5).

The Filtlong advantage in SNP calling came from recall rather than precision (Supplementary Figs S6 and S7). FPs were few for every pipeline: summed across the 13 isolates, FP SNP calls ranged from 2 to 8 at 100× and from 7 to 24 at 20× (Supplementary Fig S8). FNs separated the pipelines more clearly. At 100×, the Filtlong pipelines were the only ones with 35 or fewer FN SNP calls summed across the 13 isolates, against 55 for the raw control, and `Filtlong-Dorado` had the fewest (17, or 38 fewer than the raw control). A clear example is *E. coli* ATCC 25922202309, which had the lowest SNP F1 score of the 13 isolates for the raw control and for every non-Filtlong pipeline. At 100×, all four Filtlong pipelines produced no FNs for this isolate, compared with 26–100 FNs for the non-Filtlong pipelines. At 20×, Filtlong pipelines produced only 1–2 FNs, compared with 50–146 FNs for the non-Filtlong pipelines. However, this advantage depended on the accompanying adapter trimming tool. At 20×, `Filtlong-Barbell` produced 817 FN SNP calls, the worst result in the benchmark and about 2.5 times more than `Filtlong-Untrimmed`, which produced only 321 FNs.

Indel F1 scores for SUP-basecalled data were lower than SNP F1 scores, and the pipelines ranked differently (Fig 2B). At 100× the 17 means fell within 0.12 percentage points, from 99.40% (`Unprocessed-Barbell`) to 99.28% (`SeqKit-Porechop ABI` and `Filtlong-Barbell`). At 20× the spread widened to 0.60 percentage points, and 15 of the 17 pipelines fell below 99.00%. `SeqKit-Untrimmed` and `Filtlong-Porechop ABI` gave the highest means (both 99.05%), and the raw control gave the lowest (98.45%). The raw control missed more indels rather than calling false ones: summed across the 13 isolates, it had 90 FN and 7 FP indel calls, against 60 and 5 for `SeqKit-Untrimmed` (Supplementary Fig S8). Under HAC basecalling, the indel ranking did not follow SUP: `SeqKit-Barbell` was highest at 100× (99.16%) and `Filtlong-Dorado` at 20× (97.85%), and the raw control ranked seventh at 20× (Supplementary Fig S5).

### How accurate and contiguous are the assemblies?

At 100×, every pipeline, including the raw control, held mean assembly error rates below 1.0 error per 100 kbp for the SUP-basecalled reads (Fig 3A; HAC results in Supplementary Fig S10). The three lowest were Filtlong pipelines (`Filtlong-Dorado`, `Filtlong-Untrimmed` and `Filtlong-Porechop ABI`), at 0.22–0.25 errors per 100 kbp. The two highest were Barbell pipelines, at 0.52 (`Filtlong-Barbell`) and 0.66 (`SeqKit-Barbell`).

**Fig 3.**
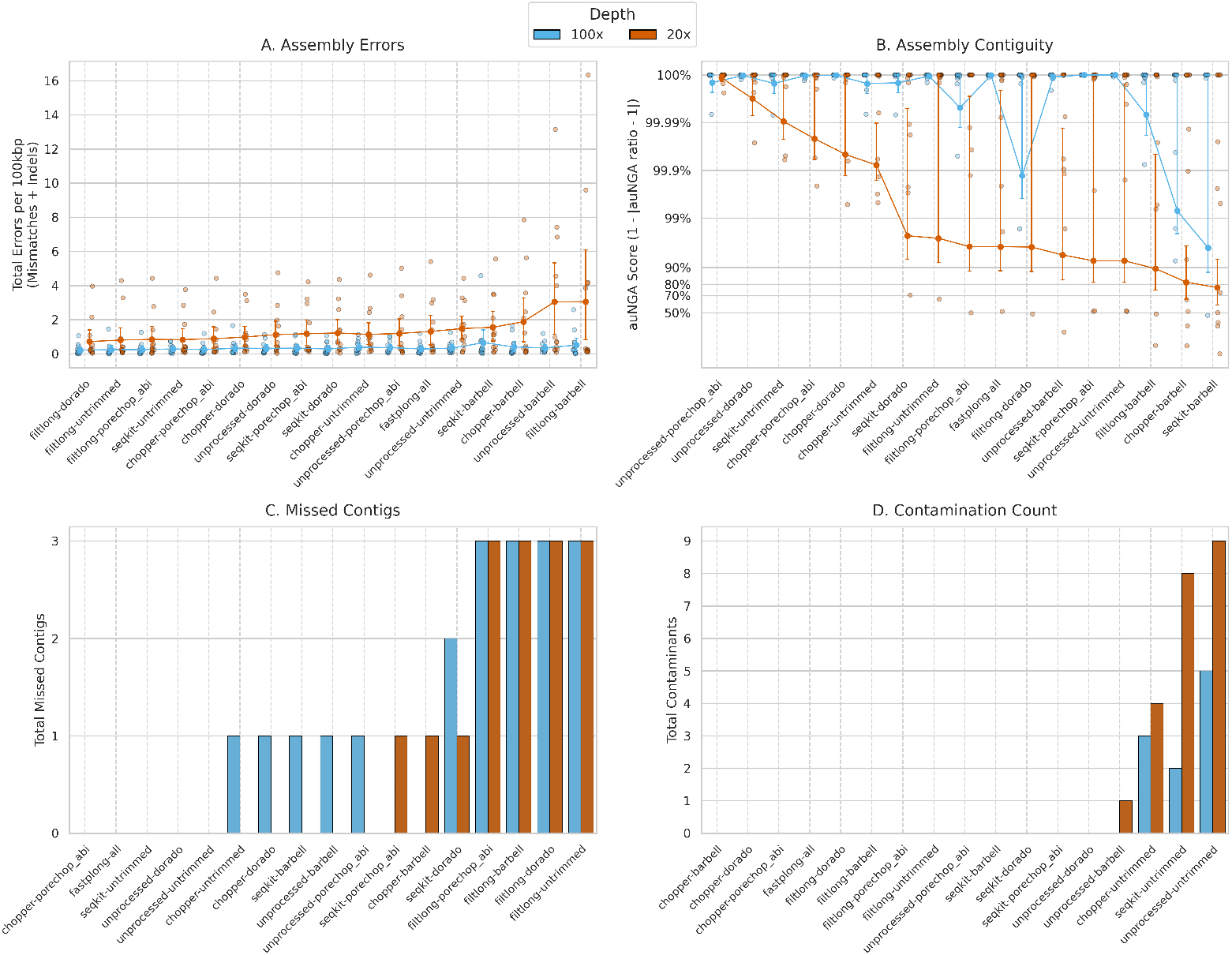
Genome assembly performance across QC pipelines and sequencing depths. Performance across four metrics at 100× (blue) and 20× (orange) depth, using SUP-basecalled reads. (A) total errors per 100 kbp (base mismatches and small indels), linear y-axis. (B) assembly contiguity: auNGA score, evaluated as 1 *−* |auNGA ratio *−* 1|, logit y-axis. (C) total count of fully and partially missed reference contigs. (D) total contaminant hits (residual adapter or barcode alignments in assembled contigs). In (A) and (B), individual points are single-sample assemblies, solid markers connected by lines are means, and error bars are 95% confidence intervals. In (C) and (D), bars are totals across the 13 isolates, on a linear y-axis.

Dropping the datasets to 20× raised error rates across all conditions and separated the pipelines by roughly a factor of four (Fig 3A). `Filtlong-Dorado` gave the lowest mean at 0.72 errors per 100 kbp, followed by `Filtlong-Untrimmed` (0.81), `SeqKit-Untrimmed` (0.83) and `Filtlong-Porechop ABI` (0.84). Barbell pipelines again produced the highest error rates: `Unprocessed-Barbell` at 3.04 and `Filtlong-Barbell` at 3.05 errors per 100 kbp. Upon further investigation, we found that the elevated Barbell error rates came from isolates prepared with the Native Barcoding Kit (Supplementary Fig S11). In the worst case, the `Filtlong-Barbell` assembly of *S. aureus* BPH2947202310 (16.34 errors per 100 kbp), the errors were clustered in regions where reads gave little or no coverage (Supplementary Fig S12 and Supplementary Section S5.1).

Contiguity differed between pipelines mainly at 20×. At 100×, most pipelines, including the raw control, held mean auNGA scores above 99.99% (Fig 3B). The only pipelines below that were `Filtlong-Dorado` (99.87%), `Chopper-Barbell` (99.30%) and `SeqKit-Barbell` (95.92%). At 20×, the three highest means, all above 99.99%, came from pipelines that applied only one of the two operations: `Unprocessed-Porechop ABI` and `Unprocessed-Dorado` trimmed adapters without quality-based processing, and `SeqKit-Untrimmed` filtered without trimming. The pipelines that applied both ranked below them. The raw control, which applied neither, held high contiguity at 100× but dropped to 92.61% at 20×. The three lowest were Barbell pipelines, at 89.57% (`Filtlong-Barbell`), 81.68% (`Chopper-Barbell`) and 77.52% (`SeqKit-Barbell`).

Under HAC basecalling all pipelines stayed above 99.98% at 100×, and at 20× the three lowest were again Barbell pipelines (Supplementary Fig S10).

### Which pipelines recover every replicon?

Every Filtlong pipeline failed to recover three reference contigs across the 13 isolates at both 100× and 20× (Fig 3C), and all three were small plasmids: one of 3,011 bp from *S. aureus* (BPH2947202310), and two of 3,173 bp and 1,919 bp from *E. coli* (ATCC 25922202309) (Supplementary Figs S13 and S14).

Among the other pipelines, missed contigs were few and changed with depth. The raw control, `Unprocessed-Dorado`, `Chopper-Porechop ABI`, `SeqKit-Untrimmed` and `Fastplong-all` recovered every replicon at both depths. The remaining eight pipelines each fully or partially missed at most two contigs at either depth. Six of them missed a contig at 100×, while five of them recovered every replicon at 20×. The raw control and `Unprocessed-Dorado` were the only pipelines to recover every replicon at both depths under both basecalling models (Supplementary Figs S13 and S14).

Some of these misses did not reproduce between runs. Repeated Flye runs changed the missed contig counts of some pipelines that kept the plasmid reads, but never for the Filtlong pipelines (Supplementary Table S3 and Supplementary Section S5.3).

Lowering Flye’s `minOverlap` parameter to 1000 also helped recover the small plasmids in non-Filtlong pipelines (Supplementary Figs S15 and S16 and Supplementary Section S5.4).

### Does trimming remove technical sequences from assemblies?

Adapter trimming was the only operation that removed residual technical sequences (adapters and barcodes) from assemblies. All three trimming tools eliminated contaminant hits whatever quality-based processing followed (Fig 3D), with one exception: `Unprocessed-Barbell` retained a single contaminant contig at 20×.

Quality-based processing on its own could not remove these sequences. The raw control carried the highest contamination counts, with five contaminant hits at 100× and nine at 20×. `Chopper-Untrimmed` retained three and four, and `SeqKit-Untrimmed` two and eight, at 100× and 20× respectively. `Filtlong-Untrimmed` was the only untrimmed pipeline with no detected contaminants at either depth. Every contaminant hit lay on a non-circular contig, within 140 bp of a contig end. No circular contig carried adapter or barcode sequence.

### Which pipeline is the best for genome assembly?

We surveyed the microbial genomics community on the importance of each of the four assembly metrics, so that the pipelines could be ranked by the priorities of the people who use these assemblies. We received 22 responses. Respondents were asked to allocate 100 points across the four metrics according to their importance. On average, they allocated 34.4 points to replicon recovery, 27.4 to sequence accuracy, 21.9 to assembly contiguity, and 16.3 to the absence of adapter and barcode contamination. Replicon recovery was therefore the most heavily weighted criterion, while the absence of adapter and barcode contamination received the lowest weight.

The composite score combines the four assembly metrics into one number. Each metric is scaled from 0 to 100, and then averaged using the survey weights. A pipeline with no sequence errors, perfect contiguity, no contaminant contigs and no missed contigs would score 100 (full definition in Materials and methods and Supplementary Section S3). Under these weights, `Chopper-Porechop ABI` scored highest at both depths, at 99.3 at 100× and 97.6 at 20× (Fig 4). It was not the best pipeline on any single metric, but it ranked well on all four: mean error rates of 0.25 and 0.89 per 100 kbp, mean auNGA scores above 99.99% at 100× and of 99.98% at 20×, no missed contigs, and no contaminant contigs. At 100×, `Fastplong-all` scored 99.2, followed by `Unprocessed-Dorado` and `SeqKit-Porechop ABI` at 99.1. At 20×, `Chopper-Dorado` scored 97.3 and `Unprocessed-Dorado` 96.9.

**Fig 4.**
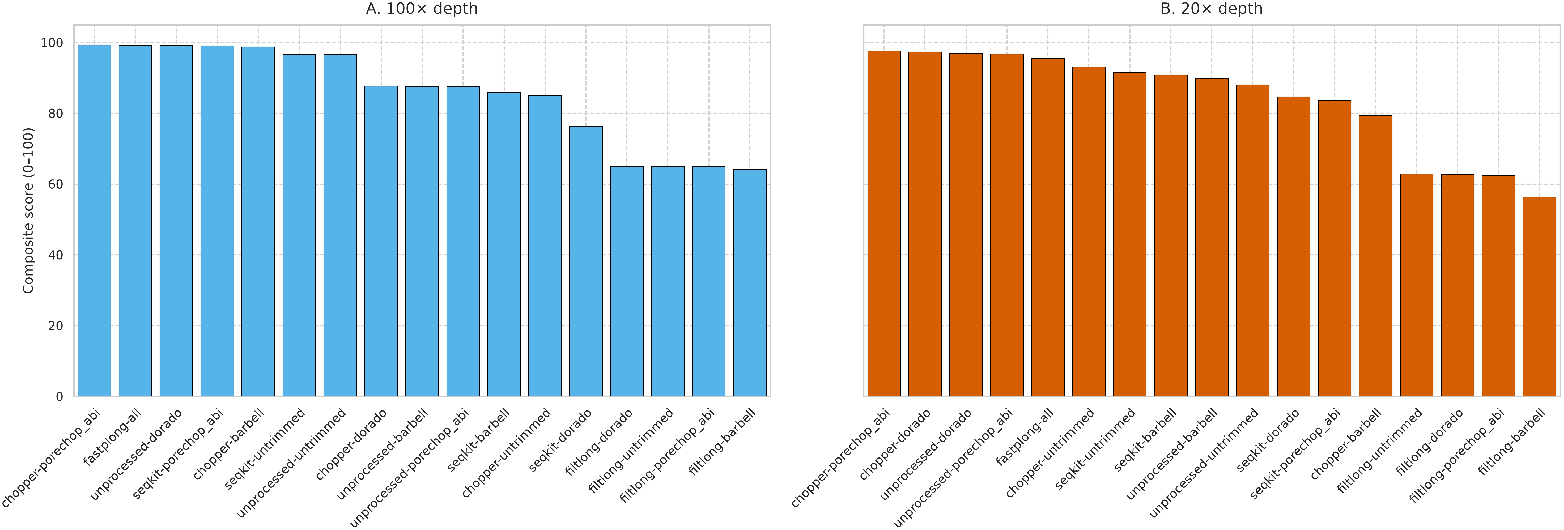
Survey-weighted composite assembly scores across QC pipelines. Assembly performance across QC tool combinations at (A) 100× and (B) 20× sequencing depths, using SUP-basecalled reads. Composite scores are a weighted arithmetic mean across four metrics using community survey weights: replicon recovery (34.4%), sequence accuracy (27.4%), assembly contiguity (21.9%), and absence of contamination (16.3%). The y-axis is the composite score on a linear 0–100 scale.

The four lowest-scoring pipelines at both depths were the four Filtlong pipelines, at 64.2–65.0 at 100× and 56.3–62.8 at 20×, despite the low sequence error rates achieved by several Filtlong pipelines. The three plasmids they failed to assemble cost them the whole replicon recovery score. With the replicon recovery weight set to zero, three Filtlong pipelines ranked first to third at 100×, led by `Filtlong-Dorado` (99.0), but all within 0.1 points of `Chopper-Porechop ABI` in fourth. At 20×, `Chopper-Porechop ABI` remained first (96.3), and the best Filtlong pipeline, `Filtlong-Untrimmed`, ranked third (95.8).

The ranking is specific to the SUP model and to these weights. Under HAC basecalling the top-ranked pipeline changed with depth, to `SeqKit-Barbell` at 100× and `Unprocessed-Porechop ABI` at 20× (Supplementary Fig S17).

## Discussion

Read preprocessing is usually treated as a routine step in bacterial genomics, applied in much the same way regardless of the downstream analysis. Our benchmark does not support that treatment. Although most of the 17 pipelines did not differ substantially from the raw, unprocessed reads for variant calling and for every assembly metric except contamination, the improvements that did appear came with trade-offs. Filtlong improved SNP F1 score for variant calling and minimised assembly error rates, but every Filtlong pipeline lost the same three small plasmids. Barbell trimming reduced contiguity and increased error rates in assemblies. Skipping adapter trimming resulted in assemblies containing residual adapter contamination. The importance of these trade-offs depends on the downstream analysis goal.

Our variant calling results agree with short-read benchmarks on adapter trimming, but diverge when applying a preprocessing strategy that combines read quality and length to score and rank reads. Bush found that adapter and quality trimming added little benefit to bacterial SNP calling pipelines built on Illumina data [14]. Adapter trimming alone behaved similarly here: trimming with Dorado, Porechop ABI, or Barbell without subsequent quality-based processing produced mean SNP F1 scores no higher than those from untrimmed reads.

This consistency reflects the behaviour of downstream alignment and variant calling tools. Aligners such as Minimap2 soft-clip read ends, preventing unmatched adapter and barcode sequences from corrupting the primary genomic alignment [16, 25].

Similarly, deep learning variant callers like Clair3 evaluate both pileup context and full-alignment representations [18], which may help them distinguish terminal non-biological artefacts from true biological variants. The difference from short reads is that ONT read lengths vary widely, which gives another property to select on. Filtlong scores and ranks reads using a weighted combination of quality and length, retaining only the highest-ranked subset up to a target base count [10]. Threshold filtering by minimum quality and length with Chopper and SeqKit did not improve F1 scores over raw reads, and neither did Filtlong using quality weight alone (Supplementary Figs S3 and S4). Filtlong’s length- and quality-weighted subset may matter most in repetitive regions. In *E. coli* ATCC 25922202309, where Filtlong pipelines had almost no FN SNP calls, Hall *et al*. traced most missed variants to two 7.5 kbp repeats with 96.5% identity, where reads from one copy mapped to the other [17]. Longer, higher-quality reads are more likely to span these repeats and map to the correct copy.

Skipping adapter trimming left residual adapter and barcode sequences in assemblies from both sequencing kits. Such sequences have previously been reported in ONT bacterial genome assemblies in public repositories [34] and can lead to taxonomic misclassification [13]. In our benchmark, contaminant sequences were located at the ends of contigs that Flye did not circularise. Flye reports a contig as circular when its path through the assembly graph closes into a loop [28], and adapter or barcode sequence at a contig end may prevent that loop from closing, leaving the replicon linear. Assemblies in which every contig circularised therefore carried no residual adapter or barcode sequences in our data. Circularisation alone does not guarantee a contaminant-free assembly: Beeloo *et al*. found a barcode on a public plasmid record that was annotated as circular [13].

Filtlong’s plasmid loss has a direct cause. Filtlong scores reads on length and quality and retains the longest, highest-quality reads up to a target base count, which biases selection against the shorter reads that come from small plasmids. The strategy lowered assembly error rates across the chromosome, but produced complete coverage dropouts for the small replicons. Under-recovery of small plasmids from ONT data is already described, but by a different mechanism. Wick *et al*. reported that plasmids below 20 kbp were underrepresented in ONT ligation datasets by a factor of roughly four, and by over 100-fold for one 2.4 kbp plasmid, whereas rapid preparations showed no comparable size bias [11]. They attribute this to library preparation: intact circular molecules that escape fragmentation have no ends for adapter ligation and are never sequenced. That mechanism does not explain our losses. Two of the three lost plasmids came from a library prepared with the Rapid Barcoding Kit, which is the preparation Wick *et al*. found to be free of size bias, and reads covering those plasmids were present in the raw dataset before preprocessing.

Projects for which plasmid recovery matters could consider using plasmid-aware assembly tools rather than a general-purpose long-read assembler. Plassembler targets plasmid assembly directly [35], and consensus approaches such as Autocycler reduce reliance on any single assembler’s behaviour [36]. For example, a Filtlong-filtered read set can give a high-accuracy chromosome in a first-pass assembly, and the plasmids can then be recovered by running Plassembler on the full, unfiltered read set. This takes longer, but it keeps the accuracy of Filtlong without losing the small plasmids.

### Choosing a pipeline

Under SUP basecalling, the Filtlong pipelines, other than `Filtlong-Barbell` at 20×, gave the highest SNP F1 scores, which makes them suitable for outbreak tracking or AMR point-mutation detection. Leaving reads untrimmed did no harm to SNP calling, although the raw control gave the lowest indel F1 score at 20×. Where indels matter and depth is low, quality-based processing is also worth applying: at 20×, `SeqKit-Untrimmed` and `Filtlong-Porechop ABI` gave the highest indel F1 scores.

For assembly under SUP basecalling, `Chopper-Porechop ABI` scored highest under the community weights at both depths, and it did so without winning any individual metric. It removed adapter sequence reliably enough to prevent contamination in assemblies, avoided the localised dropouts seen with Barbell, kept small plasmids, and held error rates low. `Unprocessed-Dorado` ranked in the top three at both depths, which makes it a reasonable option for workflows that want to avoid adding a third-party QC tool: Dorado is already in the pipeline as the basecaller. `Fastplong-all` ranked second on the composite score at 100× and, as a single tool that both trims and filters, is simple to run.

The other tools are better described by what they cost together with what they win. Filtlong gave the lowest assembly error rates in the benchmark, but lost three small plasmids in every configuration, which makes it unsuitable (on its own) for any project where replicon recovery is essential. Barbell matched the other trimmers in assembly errors on rapid-kit libraries but produced more errors on native-kit libraries at 20× (Supplementary Section S5.1), so it should be applied with the library preparation in mind. Although leaving reads untrimmed did no harm for SNP calling, it is the one option we would not recommend for assembly under any weighting: it was the only condition that produced contaminant hits, apart from one hit from `Unprocessed-Barbell` at 20×.

No single workflow fits every project, because different analyses need different read properties. To help researchers apply their own priorities rather than the survey’s, we built an interactive dashboard (https://adimascf.github.io/qcbenchnano/tool_weighting.html). Users can reweight the four assembly metrics, apply hard thresholds such as complete adapter removal or zero missed replicons, and compare isolate-level profiles across depths and basecalling models.

## Limitations

Several limitations constrain the interpretation of these results.

First, Flye does not produce deterministic results, and the missed contigs metric is affected the most. Repeated runs of Flye produced different missed contig counts for some pipelines that retained plasmid reads, and thread count also affected small plasmid recovery (Supplementary Section S5.3). All results reported here used a fixed thread count, but the metric remains sensitive to computational parameters unrelated to read quality. Absolute missed contig counts should be read as indicative rather than exact. The complete loss of all three small plasmids by every Filtlong pipeline was consistent across repeated runs, so that result does not depend on this instability.

Second, our workflow applied preprocessing *before* downsampling: quality-based processing was applied to the full-depth read set before subsampling to the target depth. This gave all datasets uniform depth before variant calling and assembly, but it also gave the length- and quality-weighted filtering tools a richer pool of candidate reads than would be available from a run that produced that little data in the first place. The Filtlong results at 20× should be read with that in mind. This approach was chosen to maintain experimental control by standardising final depth across all tool combinations, preventing yield differences from confounding downstream performance. Furthermore, it reflects typical user workflows where quality-weighted subsampling is applied to a larger raw sequencing pool to achieve a target depth. A related question remains open, because we did not assess the read sets at full depth: whether subsetting the reads with Filtlong to a lower target depth performs better than using all of them.

Third, we evaluated a single assembler. Plasmid-aware workflows and consensus assembly tools [35, 36, 37] were not used to assess the QC pipelines, and some of the replicon recovery penalties we attribute to preprocessing may be specific to Flye’s non-deterministic output.

Fourth, this benchmark assessed the impact of each pipeline but did not establish the cause of every difference. For example, we did not determine why Barbell produced more assembly errors on native-kit libraries at 20×, and we did not test why Filtlong improved SNP calling and assembly sequence accuracy. These questions are left for future work.

## Conclusion

Although subsampling reads by quality and length with Filtlong improves SNP calling in our dataset, preprocessing ONT reads for genome assembly introduces distinct trade-offs across metrics. Filtlong filtering reduces sequence errors in assemblies, but it systematically eliminates small plasmids. Similarly, skipping adapter trimming leaves residual adapter and barcode sequence in assemblies, whereas Barbell trimming reduces contiguity and increases error rates at low depth. QC preprocessing for ONT sequencing should therefore not be applied indiscriminately across all workflows; rather, tool choice must be guided by the specific downstream analytical objective.

## Supporting information

### Supplementary material

Section S1, parameter exploration for adapter trimming and quality-based processing (Supplementary Figs S1–S4, Supplementary Table S1). Section S2, assembly contiguity and the auNGA ratio. Section S3, composite score normalisation and community survey weights (Supplementary Table S2). Section S4, variant calling assessment: HAC-basecalled performance, precision and recall, and false negative and false positive counts (Supplementary Figs S5–S9). Section S5, genome assembly assessment: HAC-basecalled performance, Barbell assembly errors and library preparation kit, replicon recovery, run-to-run variation in Flye, the effect of Flye `minOverlap`, and HAC-basecalled composite ranking (Supplementary Figs S10–S17, Supplementary Table S3).

## Data availability

The reference genomes, mutated references and variant truthsets are archived on Zenodo [21]. The Snakemake workflow and the analysis and figure generation code are available at https://github.com/adimascf/qcbenchnano, and the interactive dashboard for reweighting the assembly metrics at https://adimascf.github.io/qcbenchnano/tool_weighting.html. For each isolate, we used only the POD5 files sequenced with the kit listed in Table 1; the POD5 DOIs, BioSample and reference assembly accessions for all 13 isolates are listed in `config/accessions.csv` in the code repository.

## Acknowledgements

This work was supported by resources provided by the University of Queensland Research Computing Centre’s Bunya supercomputer [38], with funding from the University of Queensland, Brisbane, Australia.

## Use of generative AI tools

Generative AI tools were used in two parts of this work. Google Gemini (`Flash 3.8`) was used mainly to build the interactive dashboard. Most of the figure generation code was also written with Gemini, starting from plotting templates written by the authors. The Snakemake workflow was designed and largely written by the authors, with Gemini helping to debug and edit parts of the code. All generated code was read, tested and corrected by the authors. Anthropic’s Claude (`Opus 5.5`) was used to revise author-written drafts of the manuscript for flow, clarity and grammar. The authors reviewed and edited every revised passage and checked it against the underlying results.

## Author contributions

**Achmad Dimas Cahyaning Furqon**: Conceptualisation, Methodology, Software, Validation, Formal analysis, Investigation, Data Curation, Writing - Original Draft, Writing - Review & Editing, Visualisation. **Ryan R Wick**: Conceptualisation, Methodology, Data Curation, Writing - Review & Editing. **Leah W Roberts**: Conceptualisation, Methodology, Writing - Review & Editing, Supervision, Funding acquisition. **Michael B Hall**: Conceptualisation, Methodology, Software, Investigation, Data Curation, Writing - Original Draft, Writing - Review & Editing, Supervision.

## Notes

https://github.com/adimascf/qcbenchnano

https://adimascf.github.io/qcbenchnano/tool_weighting.html

### Competing Interest Statement

The authors have declared no competing interest.

