## Supplement for "Evaluating Nanopore read trimming and quality filtering in bacterial genomics"

### Supplementary material for *Evaluating Nanopore read trimming and quality filtering in bacterial genomics*

#### Contents

**S1 Parameter exploration** **3**

**S2 Assembly contiguity (auNGA ratio)** **8**

**S3 Composite score normalisation** **9**

**S4 Variant calling assessment** **12**

**S5 Genome assembly assessment** **17**

**References** **26**

### S1 Parameter exploration

Several of the benchmarked tools support multiple operational modes or configurable parameters. Barbell, Porechop_ABI, Chopper and Filtlong each have alternative operational modes, and SeqKit and Fastplong take a minimum mean quality and read length. Prior to constructing the 17 benchmark pipelines, we evaluated these settings to select a single configuration for each tool. Adapter trimming and quality-based processing were evaluated separately across two independent runs. Both runs used the 13 isolates described in the main text, evaluated SUP and HAC basecalled reads, and tested downsampled depths of 100× and 20×. Settings were compared based on variant calling performance (SNP and indel F1 scores), assembly contiguity, sequence accuracy (mismatches and indels per 100 kbp), missed plasmid count, and contamination count.

#### S1.1 Adapter trimming

For adapter trimming, we compared two Barbell settings and three Porechop_ABI settings, alongside Dorado, Fastplong and untrimmed reads. No quality-based processing was applied (Figs S1 and S2). Barbell was run through `barbell kit` with default settings and with `--maximize`. Porechop_ABI was run in its default mode, which splits reads at adapters found in the middle of a read, with `--discard_middle`, which discards those reads instead, and with `--no_split`, which skips the search for middle adapters. Fastplong was run with `--disable_quality_filtering --disable_length_filtering`.

Both Barbell runs ranked last in terms of assembly errors, with the highest error rates among all tested tools and settings. However, we selected Barbell with default settings because it achieved better contiguity than the run with the `--maximize` flag, which showed a substantial drop in contiguity at 20×. For Porechop_ABI, `--discard_middle` performed better than `--no_split` on both variant calling and assembly metrics, and close to the default splitting mode. We chose `--discard_middle` because it gave higher F1 scores for both SNP and indels on SUP.

#### S1.2 Quality-based processing

For quality-based processing, we compared 11 settings across Chopper, Fastplong, Filtlong and SeqKit, with unprocessed reads as the control (Table S1, Figs S3 and S4). All settings were applied to reads trimmed by Dorado during basecalling. Fastplong was run with adapter trimming disabled, so only its filtering was tested. Quality cutoffs were Q15 for SUP and Q10 for HAC.

Under default weighting, Filtlong achieved the highest SNP F1 scores across all sequencing depths and both basecalling models, while also producing among the fewest assembly errors. Length-only weighting similarly improved SNP F1 scores at 100×, but not at 20×. Conversely, quality-only weighting resulted in the lowest F1 scores among all tested Filtlong configurations. For Chopper, `best-read-segment` extraction performed slightly better than `trim-by-quality`.

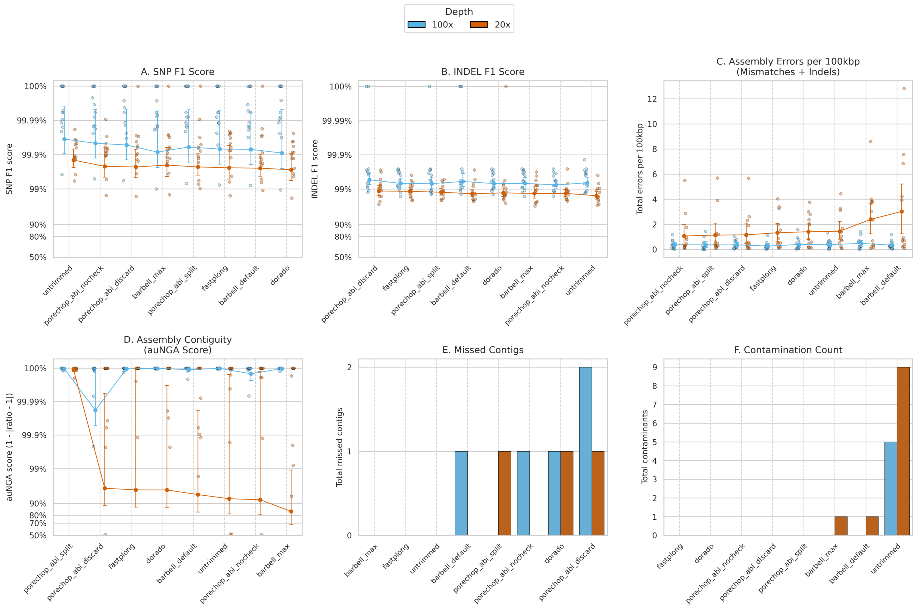

**Figure S1: Adapter trimming parameter exploration.** Barbell default against `--maximize`, and Porechop_ABI default splitting against `--discard_middle` and `--no_split`, with Dorado, Fastplong and untrimmed reads for comparison. No quality-based processing was applied. Results are shown at 100× and 20× for SUP-basecalled reads.

It achieved higher F1 scores for both SNPs and indels on SUP, produced higher contiguity, while producing assemblies with no detected contamination.

Neither length threshold performed best across all four metrics. In the SUP basecalling, raising the threshold to 1000 bp improved assembly contiguity for SeqKit and Chopper (`best-read-segment` mode), but reduced assembly completeness: it caused Chopper to miss two contigs at 20× depth and SeqKit to miss contigs at both 20× and 100×. Conversely, a 100 bp threshold recovered more replicons and reduced assembly errors, but resulted in lower contiguity and higher residual contamination for SeqKit (Figs S3). Given these opposing trade-offs, we used 100 bp as the standard minimum length for these tools in the main benchmark pipeline.

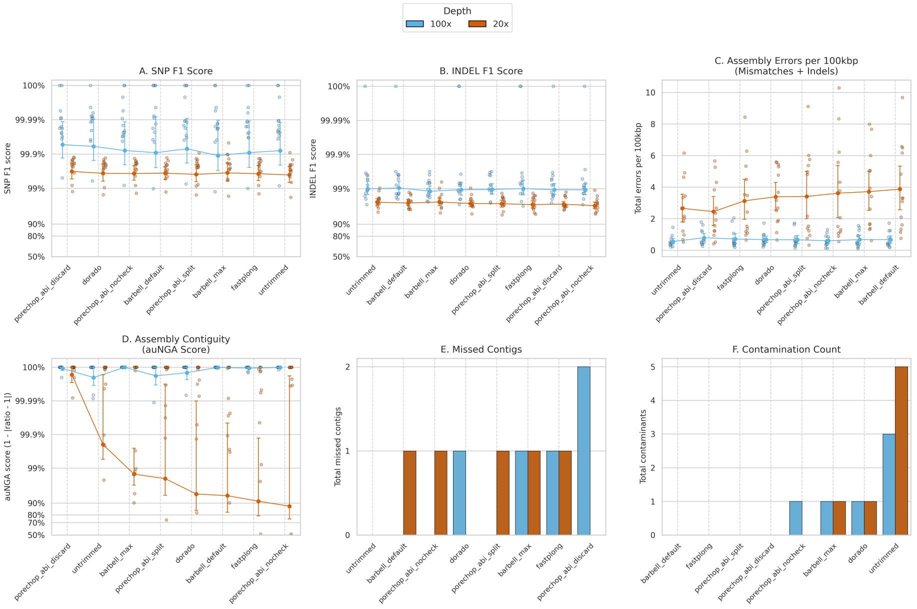

**Figure S2: Adapter trimming parameter exploration using HAC-basecalled reads.** As Fig S1, for the HAC basecalling model.

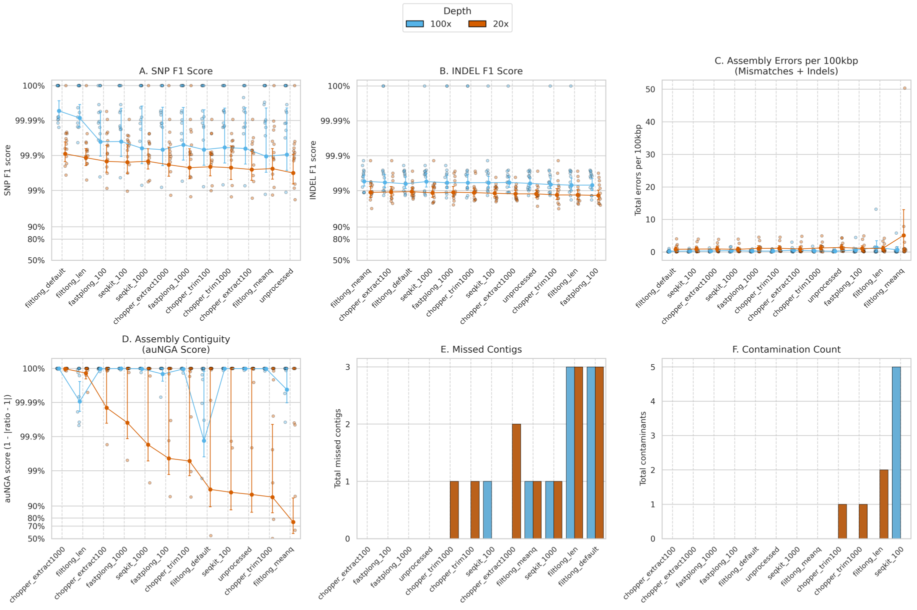

**Figure S3: Quality-based processing parameter exploration.** The 11 settings in Table S1, with unprocessed reads for comparison, applied to Dorado-trimmed reads. Results are shown at 100× and 20× for SUP-basecalled reads.

**Table S1: Quality-based processing settings explored.** $Q$ is the quality cutoff: 15 for SUP and 10 for HAC. [100×] is 100 times the genome size of each isolate. Settings marked † were used in the main benchmark. The benchmark ran Fastplong as a standalone pipeline with adapter trimming enabled, using the same 100 bp and $Q$ thresholds.

| Tool | Command | Behaviour |
|---|---|---|
| Chopper† | `chopper --trim-approach best-read-segment --cutoff` $Q$ `--minlength 100` | Extracts the longest segment with quality $\geq Q$; keeps segments $\geq$ 100 bp |
| Chopper | `chopper --trim-approach best-read-segment --cutoff` $Q$ `--minlength 1000` | As above; keeps segments $\geq$ 1000 bp |
| Chopper | `chopper --trim-approach trim-by-quality --cutoff` $Q$ `--minlength 100` | Trims read ends back to a base with quality $\geq Q$; keeps reads $\geq$ 100 bp |
| Chopper | `chopper --trim-approach trim-by-quality --cutoff` $Q$ `--minlength 1000` | As above; keeps reads $\geq$ 1000 bp |
| Fastplong | `fastplong --disable_adapter_trimming --length_required 100 --mean_qual` $Q$ | Keeps reads $\geq$ 100 bp with mean quality $\geq Q$ |
| Fastplong | `fastplong --disable_adapter_trimming --length_required 1000 --mean_qual` $Q$ | Keeps reads $\geq$ 1000 bp with mean quality $\geq Q$ |
| Filtlong† | `filtlong --target_bases` [100×] | Keeps the best reads up to the target, default weighting of length and quality |
| Filtlong | `filtlong --length_weight 0 --window_q_weight 0 --target_bases` [100×] | As above, ranking by mean quality only |
| Filtlong | `filtlong --mean_q_weight 0 --window_q_weight 0 --target_bases` [100×] | As above, ranking by length only |
| SeqKit† | `seqkit seq --min-len 100 --min-qual` $Q$ | Keeps reads $\geq$ 100 bp with mean quality $\geq Q$ |
| SeqKit | `seqkit seq --min-len 1000 --min-qual` $Q$ | Keeps reads $\geq$ 1000 bp with mean quality $\geq Q$ |

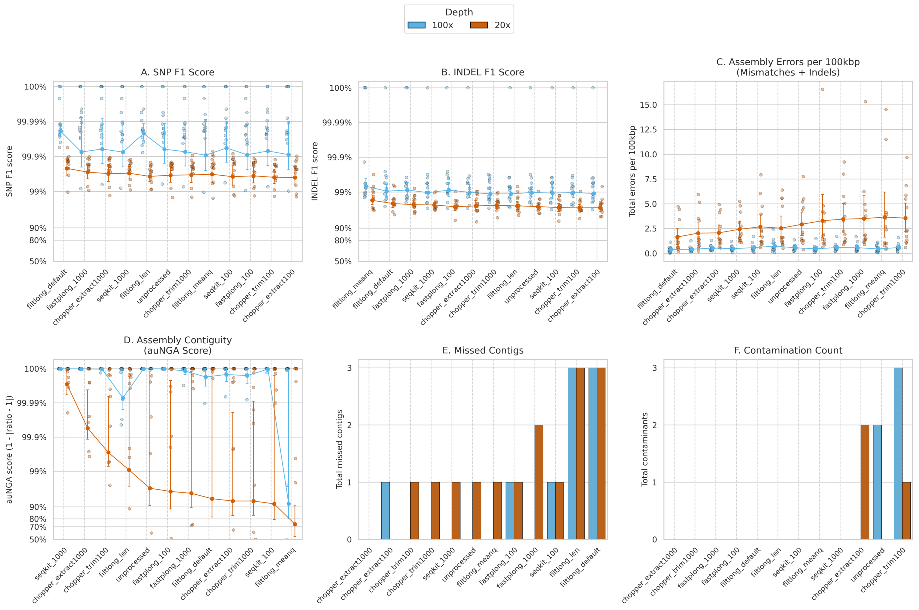

**Figure S4: Quality-based processing parameter exploration using HAC-basecalled reads.** As Fig S3, for the HAC basecalling model.

### S2 Assembly contiguity (auNGA ratio)

We assessed contiguity from the lengths of assembly segments that aligned continuously to each isolate's complete reference genome. QUAST splits contigs at detected misassemblies and excludes unaligned sequence, leaving a set of aligned blocks [1]. To calculate NGAx, these blocks are ordered from longest to shortest. NGAx is the length of the block at which their cumulative length reaches $x\%$ of the total reference length; NGA50 is the value at $x = 50$. Plotting NGAx against $x$ gives the NGAx curve. A curve that stays high indicates that much of the aligned sequence lies in long blocks, while fragmentation makes the curve fall sooner.

We summarised the whole curve by its area, auNGA, rather than using only the NGA50 point [2–4]. If $A_i$ is the length of aligned block $i$ and $G$ is the total reference length, each block contributes its length multiplied by its share of $G$:

$$\text{auNGA} = \sum_i A_i \frac{A_i}{G} = \frac{\sum_i A_i^2}{G}. \tag{S1}$$

auNGA therefore has units of length and gives more weight to long blocks. For a non-redundant assembly, it can be read as the mean block length encountered by a reference base chosen at random, with uncovered bases contributing zero. For example, splitting an intact 1 Mb alignment to a 1 Mb reference into two 0.5 Mb blocks lowers auNGA from 1 Mb to 0.5 Mb, even though the total aligned length is unchanged.

A raw auNGA value depends on the size and replicon composition of the reference. We therefore calculated a benchmark for each isolate by treating every reference replicon as one intact block. If replicon $j$ has length $R_j$, this reference value is

$$\text{auN}_{\text{ref}} = \frac{\sum_j R_j^2}{G}, \qquad G = \sum_j R_j. \tag{S2}$$

This calculation respects the separate replicons: for a reference with 3 Mb and 2 Mb replicons, the value is $(3^2 + 2^2)/(3 + 2) = 2.6$ Mb. We divided the assembly's auNGA by its own reference value to compare contiguity across isolates:

$$\text{auNGA ratio} = \frac{\text{auNGA}}{\text{auN}_{\text{ref}}} = \frac{\sum_i A_i^2}{\sum_j R_j^2}. \tag{S3}$$

An assembly that recovers each reference replicon once as a single aligned block has a ratio of 1.0. Shorter or missing aligned blocks generally lower the ratio; duplicated or redundant aligned sequence can raise it above 1.0. The ratio measures contiguity relative to the intact reference replicons, but does not identify the cause of a deviation on its own. Because both sums are dominated by the largest replicon, the loss of a small plasmid barely changes the ratio, which is why replicon recovery was scored as a separate criterion.

#### S3 Composite score normalisation

To rank the evaluated preprocessing tools for genome assembly, we aggregated the assessment metrics into a single composite score. Each metric was first normalised to a 0–100 scale.

For assembly contiguity, an auNGA ratio of $r = 1.0$ represents an assembly that recovers each reference replicon once as a single aligned block, whereas values below 1.0 indicate fragmentation and values above 1.0 indicate duplication. Deviations from 1.0 were penalised symmetrically:

$$S_{\text{contiguity}} = 100 \times \max\left(0, 1 - |r - 1.0|\right)$$

For sequence accuracy, base errors extracted from the QUAST reports were combined into total error events per 100 kbp:

$$\text{events} = \text{Mismatches per 100 kbp} + \text{Indels per 100 kbp}$$

Errors were scaled using a fixed policy anchor of 10.0 events per 100 kbp:

$$S_{\text{accuracy}} = 100 \times \max\left(0, 1 - \frac{\text{events}}{10.0}\right)$$

For residual adapter and barcode contamination, the criterion score represents the proportion of cohort isolates completely free of detected adapter and barcode hits:

$$S_{\text{contaminant}} = 100 \times \frac{N_{\text{clean}}}{13}$$

where $N_{\text{clean}}$ is the number of isolates with zero residual contaminant hits.

For replicon recovery, assemblies across the 13 isolates were evaluated using the sum of fully and partially missed contigs ($M = \text{full\_missed} + \text{partial\_missed}$) with a denominator of 3.0:

$$S_{\text{replicon}} = 100 \times \max\left(0, 1 - \frac{M}{3.0}\right)$$

An assembly pipeline retaining all replicons receives a score of 100, while losing three or more replicons across the cohort reduces $S_{\text{replicon}}$ to 0.

Fixed policy anchors were adopted instead of dynamic min–max normalisation to avoid exaggerating minor performance differences. At 100× under SUP, mean error rates ranged only from 0.22 to 0.66 errors per 100 kbp. Min–max scaling would turn this 0.44 difference into a 100-point gap, whereas the 10.0-event anchor keeps the mean accuracy score of every pipeline above 93. For replicon recovery, a denominator of 3.0 reflects the three small plasmids present in the benchmark cohort, defining a score of zero as the complete loss of all small plasmids across the isolates.

For contiguity ($S_{\text{contiguity}}$) and sequence accuracy ($S_{\text{accuracy}}$), scores were calculated for each isolate individually before taking the cohort arithmetic mean ($C_{\text{contiguity}}$ and $C_{\text{accuracy}}$). Contaminant removal ($C_{\text{contaminant}} = S_{\text{contaminant}}$)

and replicon recovery ($C_{\text{replicon}} = S_{\text{replicon}}$) were evaluated across the complete cohort.

To set the weights, we asked the microbial genomics community, through a microbial genomics Slack workspace, to complete a survey on the importance of the four assembly metrics, and we received 22 responses. Each respondent distributed 100 points across the four metrics (Table S2). The mean of these responses provided the weights for the composite score:

$$S = \frac{w_{\text{acc}} C_{\text{accuracy}} + w_{\text{cont}} C_{\text{contiguity}} + w_{\text{contam}} C_{\text{contaminant}} + w_{\text{rep}} C_{\text{replicon}}}{100}$$

**Table S2: Community survey weights.** Point allocations across the four assembly criteria from the 22 survey responses. Each respondent distributed 100 points. Respondents are numbered in order of submission. Means were rounded to one decimal place so that the four weights sum to 100 (largest-remainder rounding). The mean of the responses was used as the weights for the composite score.

| Respondent | Sequence accuracy | Contiguity | Absence of contamination | Replicon recovery |
|---|---|---|---|---|
| R1 | 40 | 35 | 5 | 20 |
| R2 | 30 | 20 | 35 | 15 |
| R3 | 20 | 20 | 25 | 35 |
| R4 | 25 | 15 | 10 | 50 |
| R5 | 45 | 5 | 5 | 45 |
| R6 | 10 | 20 | 35 | 35 |
| R7 | 25 | 15 | 10 | 50 |
| R8 | 25 | 40 | 5 | 30 |
| R9 | 35 | 10 | 25 | 30 |
| R10 | 0 | 25 | 25 | 50 |
| R11 | 30 | 25 | 20 | 25 |
| R12 | 40 | 10 | 10 | 40 |
| R13 | 25 | 25 | 25 | 25 |
| R14 | 39 | 17 | 13 | 31 |
| R15 | 50 | 10 | 20 | 20 |
| R16 | 10 | 15 | 25 | 50 |
| R17 | 25 | 25 | 10 | 40 |
| R18 | 20 | 40 | 15 | 25 |
| R19 | 30 | 10 | 10 | 50 |
| R20 | 35 | 20 | 10 | 35 |
| R21 | 35 | 20 | 15 | 30 |
| R22 | 10 | 60 | 5 | 25 |
| Mean | 27.4 | 21.9 | 16.3 | 34.4 |
| SD | 12.5 | 12.6 | 9.4 | 11.1 |
| Range | 0–50 | 5–60 | 5–35 | 15–50 |

### S4 Variant calling assessment

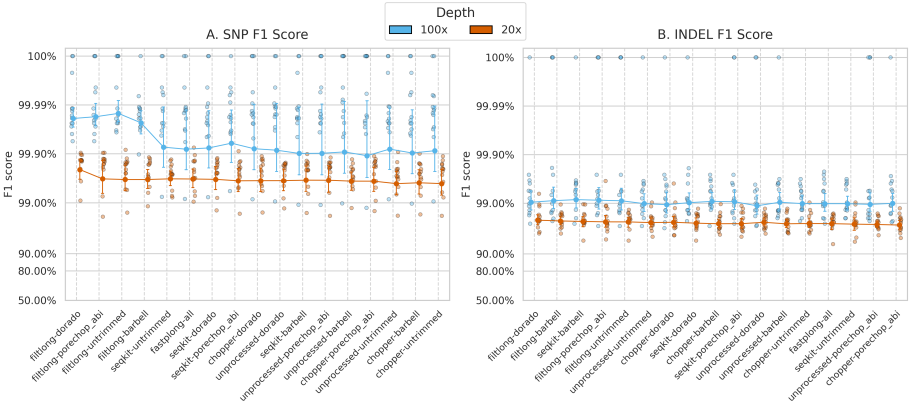

**Figure S5: Variant calling performance using HAC-basecalled reads.** As Fig 2 in the main text, for the HAC basecalling model. F1 scores for (A) SNP and (B) indel variant calling at 100× (blue) and 20× (orange). The y-axis is on a logit scale. Individual points are single-sample runs, solid markers are mean F1 scores, and error bars are 95% confidence intervals.

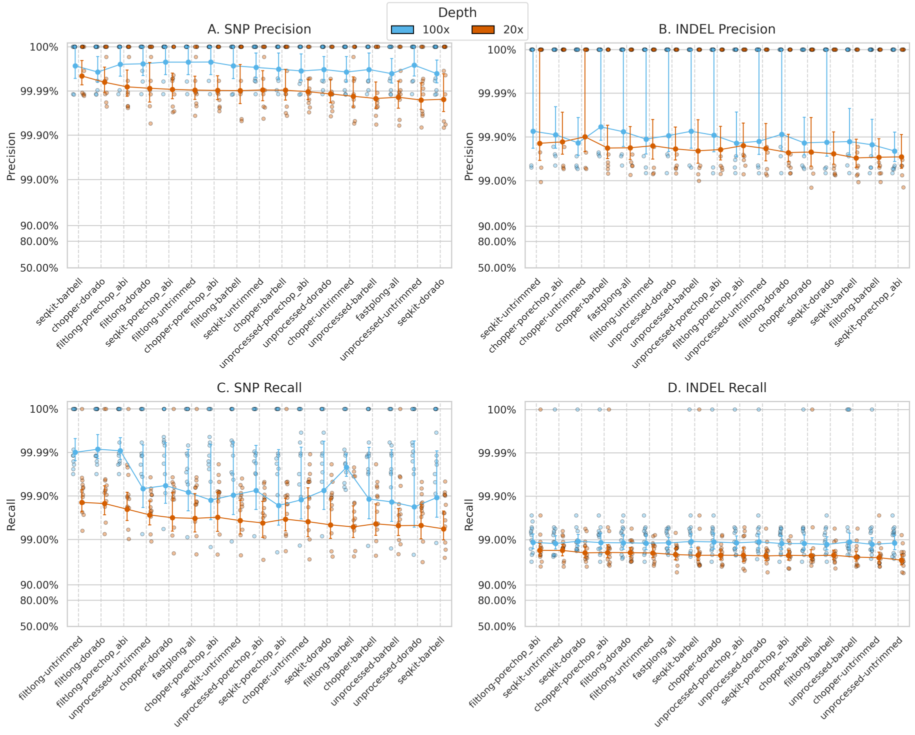

**Figure S6: Variant calling precision and recall across QC pipelines.** (A) SNP precision, (B) indel precision, (C) SNP recall and (D) indel recall across all 17 pipelines at 100× (blue) and 20× (orange), using SUP-basecalled reads. The y-axis is on a logit scale. Individual points are single-sample runs, solid markers are means, and error bars are 95% confidence intervals.

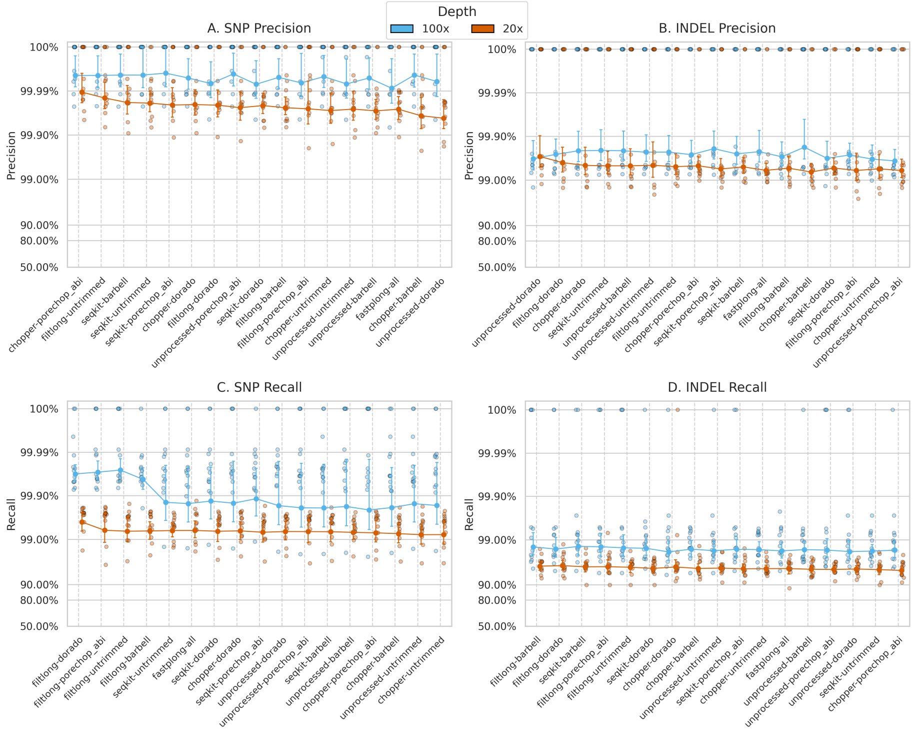

**Figure S7: Variant calling precision and recall using HAC-basecalled reads.** As Fig S6, for the HAC basecalling model.

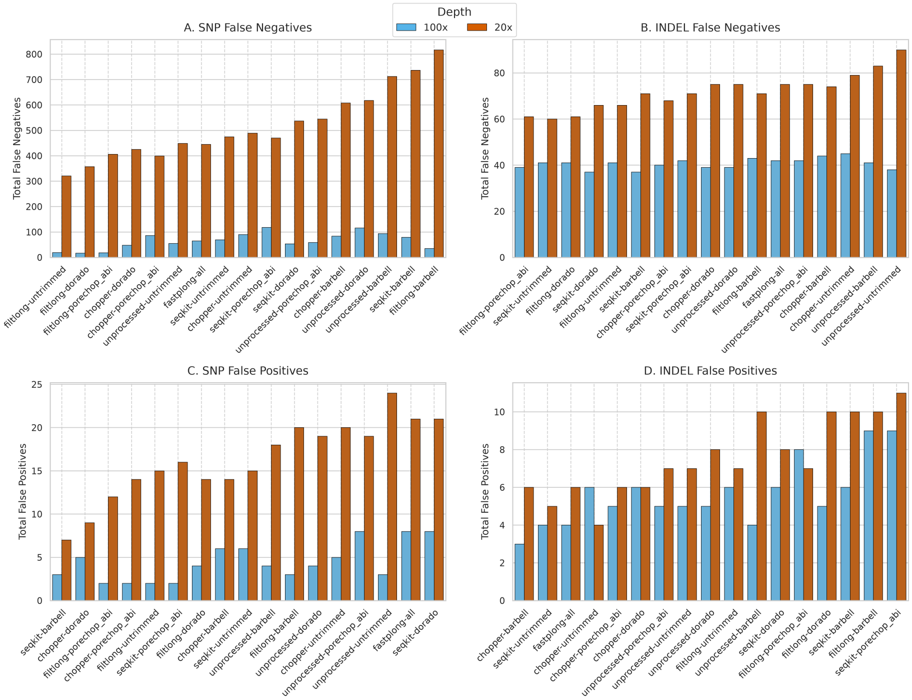

**Figure S8: Variant calling false negative and false positive counts across QC pipelines.** Total false negatives for (A) SNPs and (B) indels, and total false positives for (C) SNPs and (D) indels, across the 17 pipelines at 100× (blue) and 20× (orange) depth, using SUP-basecalled reads. Bars are counts summed across all 13 isolates, on a linear y-axis whose scale differs between panels. Within each panel, pipelines are ordered from the fewest to the most errors summed over both depths, so the order differs between panels.

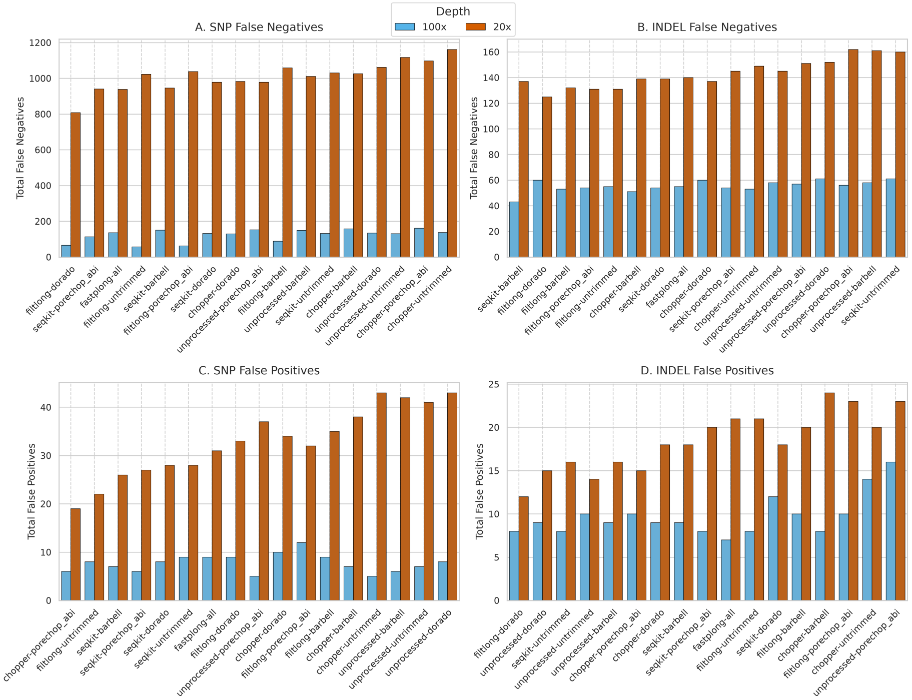

**Figure S9: Variant calling false negative and false positive counts using HAC-basecalled reads.** As Fig S8, for the HAC basecalling model.

### S5 Genome assembly assessment

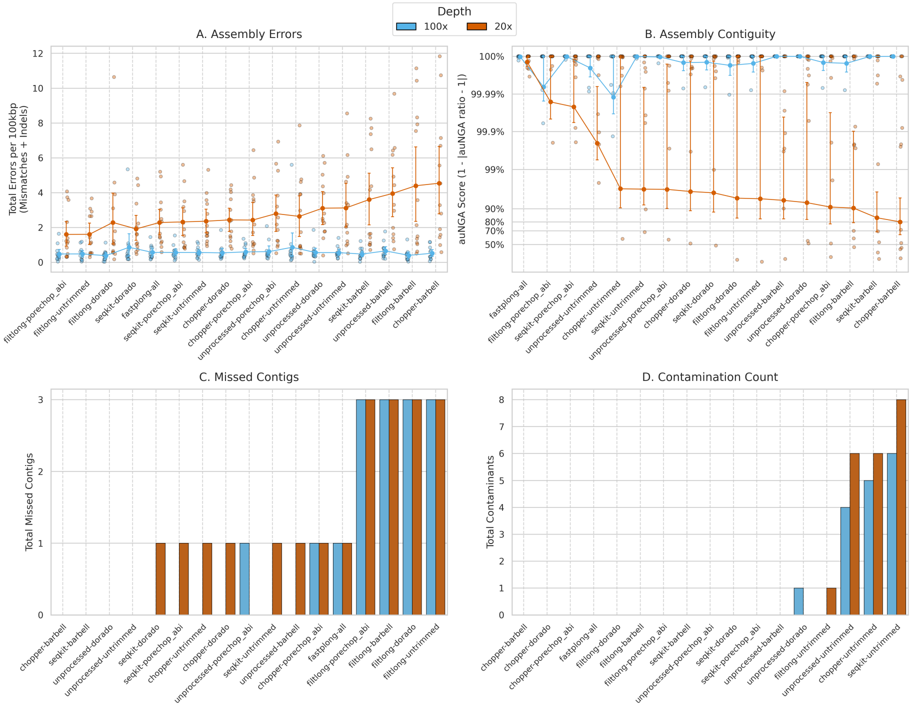

**Figure S10: Genome assembly performance using HAC-basecalled reads.** As Fig 3 in the main text, for the HAC basecalling model. (A) Total errors per 100 kbp (linear y-axis), (B) auNGA score (logit y-axis), (C) total missed contigs and (D) contaminant contig hits (linear counts summed across the 13 isolates), at 100× (blue) and 20× (orange).

#### S5.1 Barbell assembly errors and library preparation kit

At 20×, the Barbell pipelines had the highest assembly error rates, although these errors were not evenly distributed across the cohort (Fig S11). Isolates prepared with the Native Barcoding Kit had higher mean error rates than rapid-kit isolates with every pipeline, but the difference was largest for Barbell. At 20×, the four Barbell pipelines averaged 2.28–5.50 errors per 100 kbp on native-kit isolates, against 1.12–1.65 for every other pipeline. On rapid-kit isolates, the Barbell pipelines (0.20–0.70) fell within the range of the other pipelines (0.20–1.43).

We examined the highest-error case, the `Filtlong-Barbell` assembly of *S. aureus* BPH2947_202310 at 20×, which had 16.34 errors per 100 kbp. Its mismatches and indels were clustered at specific positions along the genome (Fig S12A). Mapping the `Filtlong-Barbell` reads for that isolate to the reference genome showed that these positions corresponded to regions with little

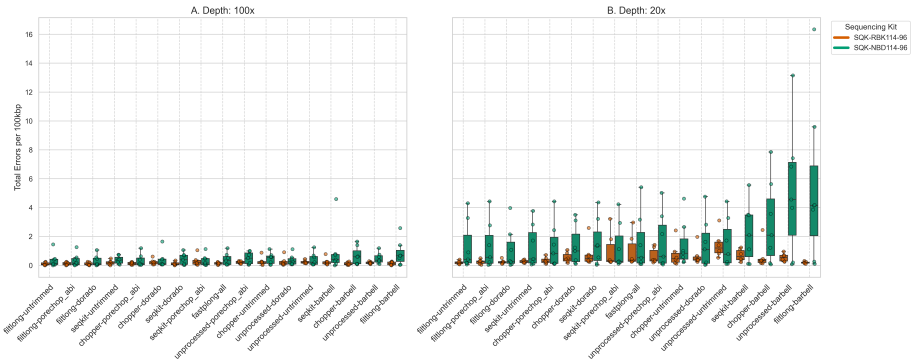

**Figure S11: Error rate in native barcoding kit (SQK-NBD114-96) vs rapid barcoding kit (SQK-RBK114-96).** Total errors per 100 kbp (base mismatches and small indels; linear y-axis) at (A) 100× and (B) 20×, using SUP-basecalled reads. Colors represent the sequencing kit used for each isolate.

to no read coverage (Fig S12B). In contrast, the `Filtlong-Untrimmed` reads, which gave a low-error assembly, covered this region well. These findings suggest that the elevated error rate was associated with insufficient read coverage in these regions.

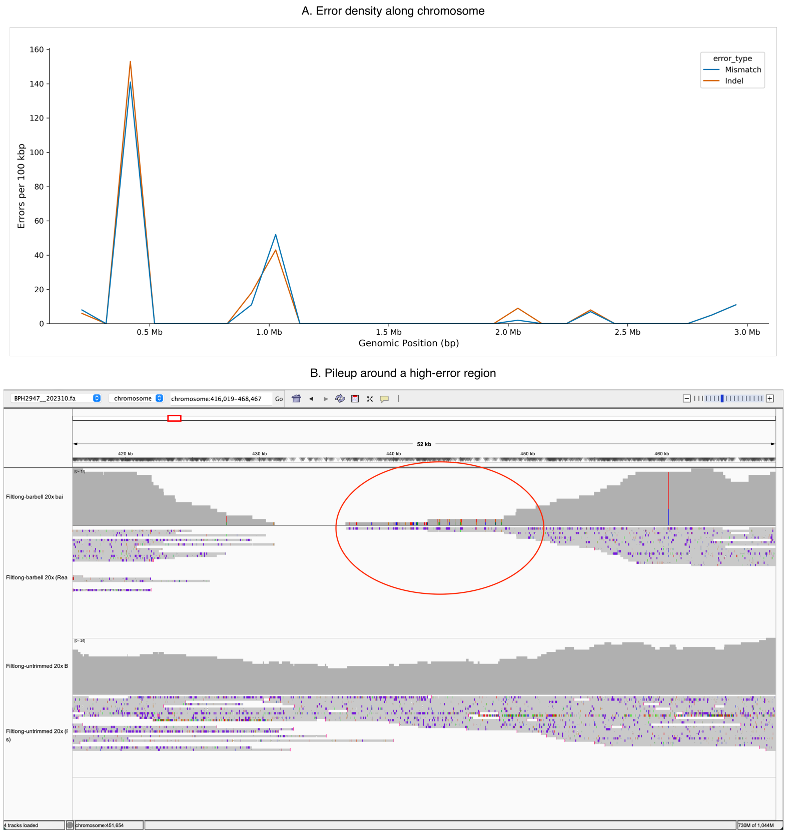

**Figure S12: High error regions from a `Filtlong-Barbell` assembly.** *S. aureus* BPH2947__202310 at 20× sequencing depth. (A) Error density distribution, with colors representing mismatches (blue) and indels (orange). (B) An IGV screenshot of the high-error region (416–468 kbp), showing the `Filtlong-Barbell` reads (top track) and the `Filtlong-Untrimmed` reads (bottom track) mapped to the reference genome.

#### S5.2 Replicon recovery

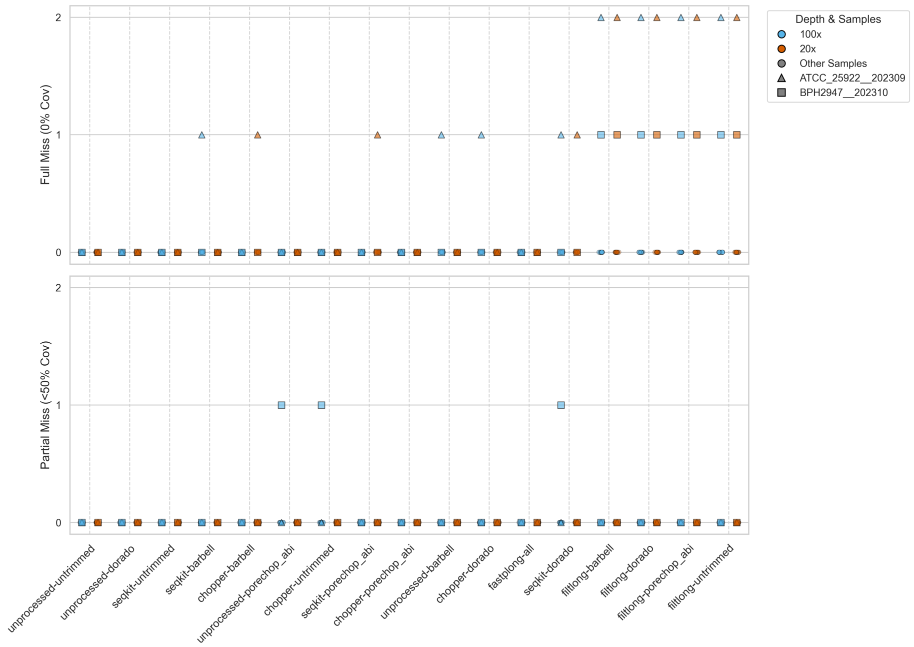

**Figure S13: Frequency of missed plasmids across QC pipelines.** Counts of full (0% coverage, top) and partial (< 50% coverage, bottom) plasmid misses across the 17 pipelines, using SUP-basecalled reads. Colors represent sequencing depths (100× in blue, 20× in orange), and point shapes denote individual bacterial isolates.

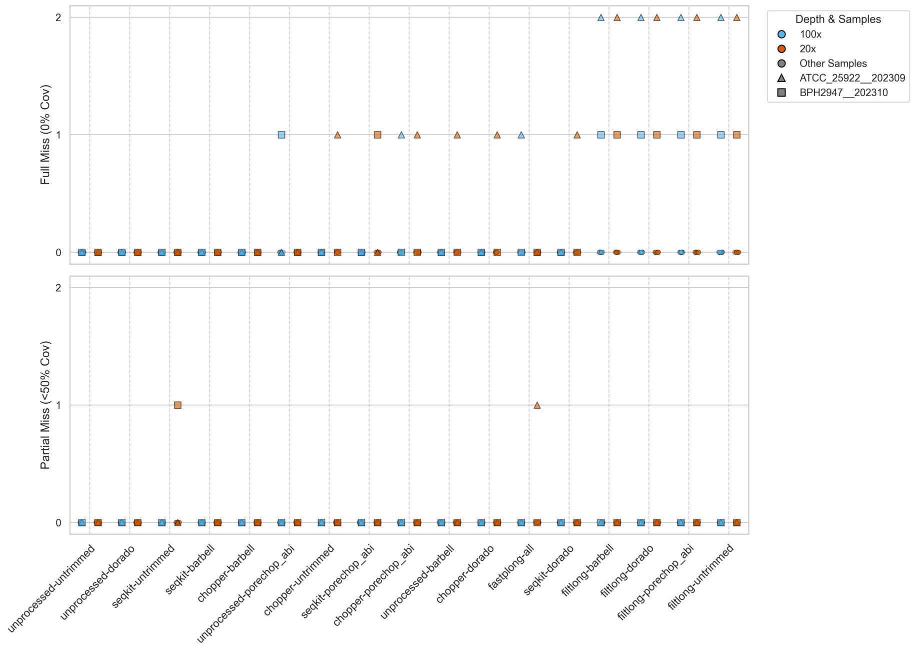

**Figure S14: Frequency of missed plasmids using HAC-basecalled reads.** As Fig S13, for the HAC basecalling model.

#### S5.3 Run-to-run variation in Flye

Small plasmid recovery by Flye was not fully reproducible between runs. Flye generates disjointigs in parallel, and this step can give different output depending on how threads are scheduled (Flye issue #509, `https://github.com/mikolmogorov/Flye/issues/509`).

To test the effect of thread count, we assembled the `SeqKit-Dorado` reads of *S. aureus* BPH2947__202310 with 1, 2, 4, 8, 16 and 32 threads, four runs each (Table S3). In the benchmark, this read set only partially assembled the 3,011 bp plasmid at 100×. At 100×, recovery depended on thread count. With one thread the plasmid was fully assembled in all four runs. With 4 or 16 threads it was only partly assembled (38.2–38.4%) in every run, and with 2, 8 or 32 threads the result changed between runs. At 20×, the plasmid was 99.4% covered at every thread count.

All benchmark assemblies used 16 threads. Across the stored reruns of the benchmark, the missed contig counts of the Filtlong pipelines never changed. The run-to-run variation affected only pipelines that kept the plasmid reads.

**Table S3: Effect of Flye thread count on small plasmid recovery.** Coverage (%) of the 3,011 bp plasmid of *S. aureus* BPH2947__202310 in Flye assemblies of the `SeqKit-Dorado` SUP reads, in four runs at each thread count.

| Threads | 100× (runs 1–4) | 20× (runs 1–4) |
|---|---|---|
| 1 | 100, 100, 100, 100 | 99.4, 99.4, 99.4, 99.4 |
| 2 | 100, 39.1, 100, 100 | 99.4, 99.4, 99.4, 99.4 |
| 4 | 38.2, 38.2, 38.2, 38.2 | 99.4, 99.4, 99.4, 99.4 |
| 8 | 38.4, 38.4, 38.4, 100 | 99.4, 99.4, 99.4, 99.4 |
| 16 | 38.4, 38.4, 38.2, 38.2 | 99.4, 99.4, 99.4, 99.4 |
| 32 | 38.2, 100, 50.1, 38.4 | 99.4, 99.4, 99.4, 99.4 |

#### S5.4 Effect of Flye `minOverlap`

By default, Flye chooses its minimum overlap between reads (`minOverlap`) from the read length N90 [5]. In a bacterial ONT dataset, abundant chromosomal reads push the N90 above the length of small plasmids, so the assembler cannot find the overlaps it needs to assemble the plasmid reads. Setting `minOverlap` to 1000 allowed Flye to use shorter overlaps and recovered small plasmids in the non-Filtlong pipelines (Figs S15 and S16).

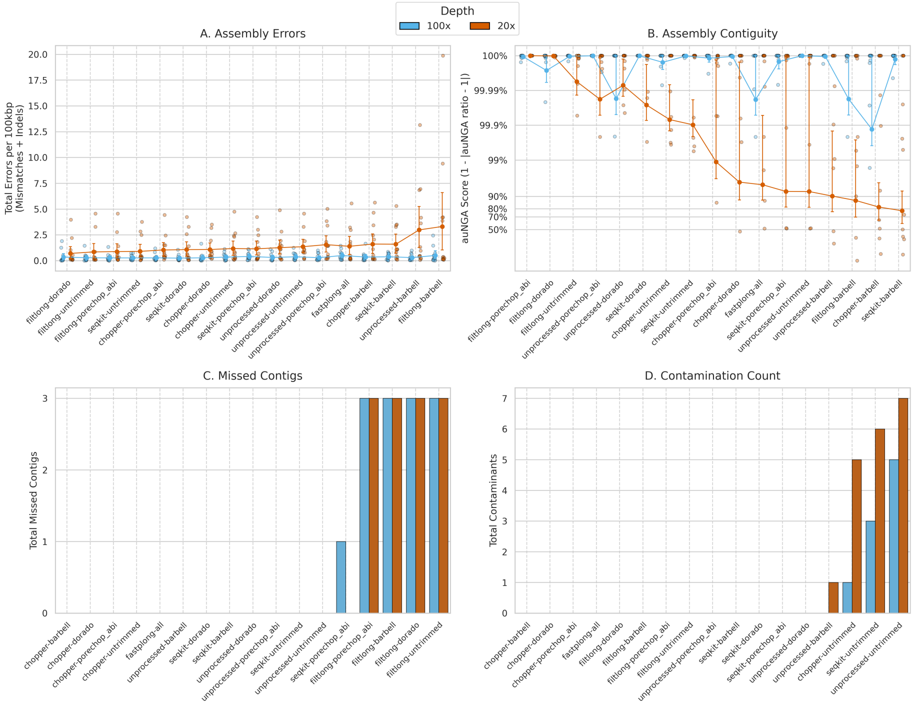

**Figure S15: Effect of Flye `minOverlap` on small plasmid recovery.** Missed contig counts under `minOverlap` set to 1000, with the corresponding effect on contiguity, mismatches, indels and contamination count, for SUP-basecalled reads.

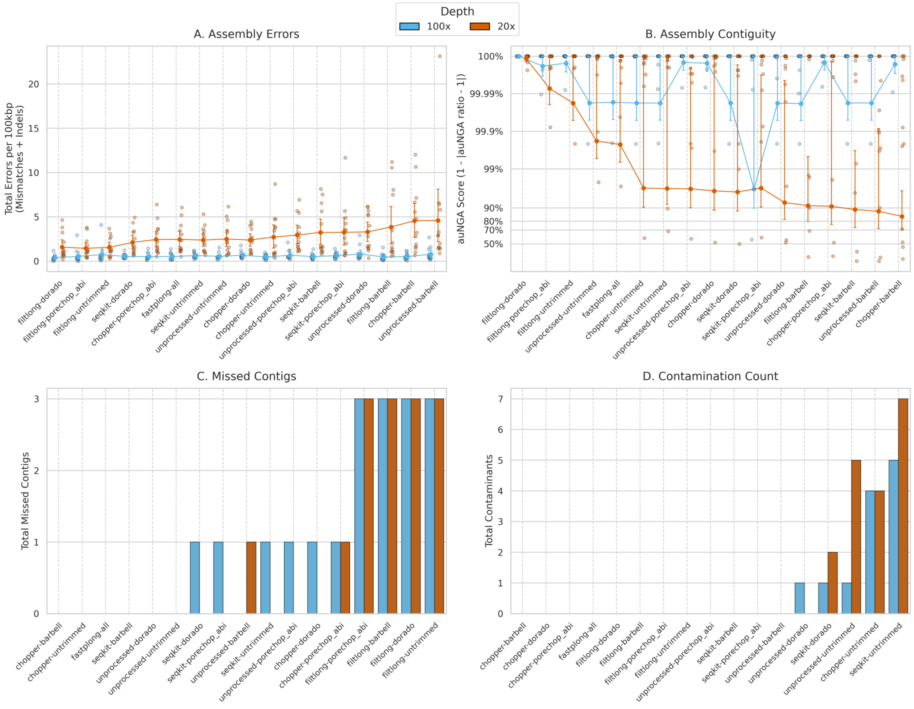

**Figure S16: Effect of Flye `minOverlap` on small plasmid recovery using HAC-basecalled reads.** As Fig S15, for the HAC basecalling model.

#### S5.5 Composite ranking

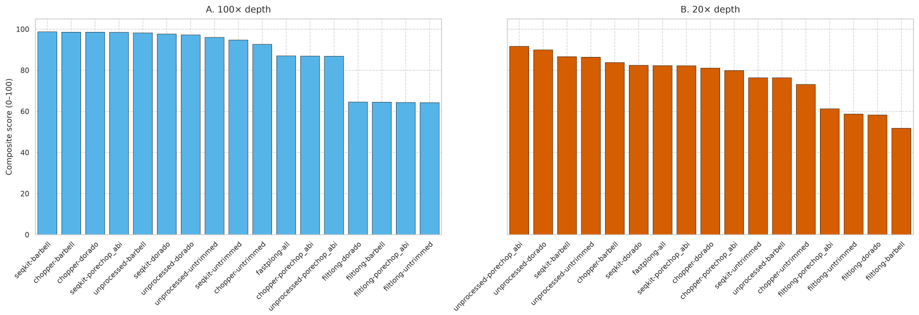

**Figure S17: Survey-weighted composite assembly scores using HAC-basecalled reads.** As Fig 4 in the main text, for the HAC basecalling model, at (A) 100× and (B) 20×. The y-axis is the composite score on a linear 0–100 scale.
